# Structure-based identification and functional characterization of an essential lipocalin in the malaria parasite *Plasmodium falciparum*

**DOI:** 10.1101/2020.02.13.947507

**Authors:** Paul-Christian Burda, Thomas Crosskey, Katharina Lauk, Aimo Zurborg, Christoph Söhnchen, Benjamin Liffner, Louisa Wilcke, Jan Strauss, Cy Jeffries, Dmitri I. Svergun, Danny W. Wilson, Matthias Wilmanns, Tim-Wolf Gilberger

## Abstract

Proteins of the lipocalin family are known to bind small hydrophobic ligands and are involved in various physiological processes ranging from lipid transport to oxidative stress responses. The genome of the malaria parasite *Plasmodium falciparum* contains a single protein PF3D7_0925900 with a lipocalin signature. Using crystallography and small-angle X-ray scattering, we show that the protein has a tetrameric structure of typical lipocalin monomers, hence we name it *P. falciparum* lipocalin (*Pf*LCN), the first lipocalin structurally and functionally characterized in a single-celled eukaryote. We show that *Pf*LCN is expressed in the intraerythrocytic stages of the parasite and localizes to the parasitophorous and food vacuoles. Conditional knockdown of *Pf*LCN impairs parasite development, which can be rescued by treatment with the radical scavenger Trolox or by temporal inhibition of hemoglobin digestion. This suggests a key function of *Pf*LCN in counteracting oxidative stress induced cell damage during multiplication of parasites within red blood cells.

## INTRODUCTION

Lipocalins are a diverse family of small proteins (typically 160 – 180 residues) whose functional hallmark is their binding to small hydrophobic molecules. Although lipocalins show low overall primary sequence conservation, they share a similar β-barrel structure consisting of eight β-strands that enclose a hydrophobic binding pocket and all are part of the larger protein superfamily of calycins (reviewed in (Flower *et al*, 2000)). The conservation of particular residues and interactions across the lipocalin protein family supports a common evolutionary origin and lipocalins have been identified and characterized in mammals, insects, plants and bacteria, with key roles in specialized biological functions including the transport of fatty acids, lipids and hormones, control of cell regulation, cryptic coloration, enzymatic synthesis of prostaglandins, and protection from oxidative stress (Flower, 1996; Ganfornina *et al*, 2008; Sanchez *et al*, 2006; Walker *et al*, 2006; Charron *et al*, 2008; Bishop, 2000). Yet, lipocalins have not been identified in archaea, and their identity and function in unicellular eukaryotes, including important human parasites, remains poorly understood.

The malaria parasite *Plasmodium falciparum*, which is responsible for more than 400,000 deaths each year (WHO, 2019), is an intracellular parasite that proliferates within a vacuole in its host cell. It establishes itself in its vertebrate host by using hepatocytes for an initial multiplication step before exploiting erythrocytes for exponential growth. To fuel the parasite’s rapid growth over its 48 hour erythrocytic lifecycle, the parasite not only feeds on its host cell but also transforms it, establishing new transport systems that allow direct access to nutrients from the surrounding blood stream. Underpinning the malaria parasite’s rapid growth are highly efficient membrane synthesis and recycling pathways that provide the structural constituents for cell growth and multiplication. For instance, the phospholipid content of the erythrocyte increases almost fivefold during intraerythrocytic development, as the parasite generates the membranes needed for growth and division (Tran *et al*, 2016). Fatty acids, the building blocks of lipids, are typically scavenged from the host but the presence a functional FASII system in the apicoplast (specialized chloroplast-derived organelle) also enables the parasite to synthesize fatty acids *de novo* (reviewed in (Tarun *et al*, 2009)). How these essential building blocks of membranes are transported from the host and within the parasite is not known.

The rapid growth of the parasite in its host cell not only depends on sufficient nutritional uptake but also on a tightly controlled redox homeostasis. In erythrocytes, the primary amino acid source for the parasite is host cell hemoglobin (Hb). Hb degradation occurs in the acidic environment of the digestive or food vacuole (FV) and involves the endocytic uptake of host cell cytosol across both the parasitophorous vacuole membrane (PVM) and the parasite plasma membrane (PPM) (reviewed in (Wunderlich *et al*, 2012)). A major waste product of Hb digestion is toxic free heme (ferriprotoporphyrin IX), most of which is detoxified by biomineralization into the inert crystalline form hemozoin. Oxidation of Hb-bound iron during Hb digestion leads to the release of superoxide radicals. The superoxide anion dismutates into hydrogen peroxide, which together with the free iron of Hb digestion can lead to the formation of hydroxyl radicals via the Fenton reaction. These radicals are highly reactive and can cause membrane damage by lipid peroxidation (Jortzik & Becker, 2012; Atamna & Ginsburg, 1993). It is thus not surprising that *Plasmodium* infected erythrocytes show much higher levels of reactive oxygen species (ROS) relative to uninfected erythrocytes (Atamna & Ginsburg, 1993). Protection against oxidative damage is therefore of particular importance for parasite survival and parasites remain reliant on various detoxification pathways including glutathione and thioredoxin redox systems, members of which have been shown to be essential for blood stage parasite development (reviewed in (Müller, 2004; Jortzik & Becker, 2012)).

In bacterial and eukaryotic cells, members of the lipocalin family are key components of lipid transport and can also mediate control of ROS damage to membranes (Flower, 1996; Ganfornina *et al*, 2008; Sanchez *et al*, 2006; Walker *et al*, 2006; Charron *et al*, 2008; Bishop, 2000). To date, no lipocalin has been structurally or functionally characterized in the diverse parasitic phylum of Apicomplexa, and a potential role for members of this protein family in survival and virulence of these important human and livestock pathogens has not been investigated. Here, we identify and characterize a new lipocalin in the human malaria parasite *Plasmodium falciparum* (PlasmoDB: PF3D7_0925900) based on structural analysis and show that it is essential for parasite survival within erythrocytes.

## RESULTS

### Identification and evolutionary analysis of lipocalin-like proteins in *P. falciparum* and related protists

Database analysis on PlasmoDB (Aurrecoechea *et al*, 2009) revealed PF3D7_0925900 as the only gene in the genome of *P. falciparum* 3D7 containing a calycin and lipocalin superfamily signature (Mitchell *et al*, 2019; Pandurangan *et al*, 2019). The encoded protein of 217 amino acids length, which we term *P. falciparum* lipocalin (*Pf*LCN), has an unknown function. It contains an N-terminal signal peptide and its mRNA is most highly expressed during trophozoite development (Otto *et al*, 2010). Sequence comparisons show that it is highly conserved between different *Plasmodium* species with 65% to 99.5% pairwise sequence identities (Figure S1). In contrast, low pairwise identities of 23% to 27% were observed for lipocalin-like proteins in *Toxoplasma gondii* (GenBank CEL71535), *Neospora caninum* (GenBank XP_003879719) and *Hammondia hammondi* (GenBank KEP62611), and blastp searches did not yield any significant sequence alignments to species outside the apicomplexan lineage, where overall pairwise sequence identities to known lipocalin proteins dropped below 20%, making it difficult to detect more distant homologous proteins at the sequence level. Therefore, we relied upon annotations based on libraries of protein signatures provided by the SUPERFAMILY and other databases integrated in InterPro to identify additional putative lipocalins (Mitchell *et al*, 2019; Pandurangan *et al*, 2019). Interestingly, when we did a comparative analysis of the genomes of other apicomplexan parasites and their closest free-living non-parasitic relatives for the presence of proteins belonging to the lipocalin (SSF50814 SUPERFAMILY identifier) and calycin (IPR012674, InterPro identifier) homologous superfamily, we found that most parasitic species encode one to three putative lipocalin-like genes only, while in most of the free living relatives a substantially higher number of putative lipocalin-like genes can be found even when different genome sizes are taken into account (Figure S2). This may hint to a potential gene reduction of the lipocalin family during apicomplexan evolution as an adaptation to the intracellular parasitic lifestyle.

### X-ray crystallography of recombinant *Pf*LCN reveals a typical lipocalin structure

Lipocalins have a characteristic structure consisting of eight anti-parallel β-barrel strands that form a hydrophobic binding pocket (Flower, 1996). To validate *Pf*LCN as a lipocalin, we recombinantly expressed it in *E. coli* without the signal peptide (all residue numbering and molecular weights that follow reflect this) and determined its structure using X-ray crystallography to a resolution of 2.85 Å (Figures 1A and B, Figure S3). The structure has been deposited with the PDB ID 6TLB. In this crystal structure, *Pf*LCN is arranged as a tetramer and the monomers superimpose with a root mean square deviation (RMSD) <0.5 Å over all Cα atoms, indicating high similarity (Figure 1A, Movie S1). The tetramer displays a two-fold symmetry and is composed of a dimer of dimers. The dimeric interface is very stable, with a solvation energy of around −17 kcal mol^−1^ calculated using the PISA server (Krissinel & Henrick, 2007), and this is strongly driven by hydrophobic effects.

**Figure 1.**
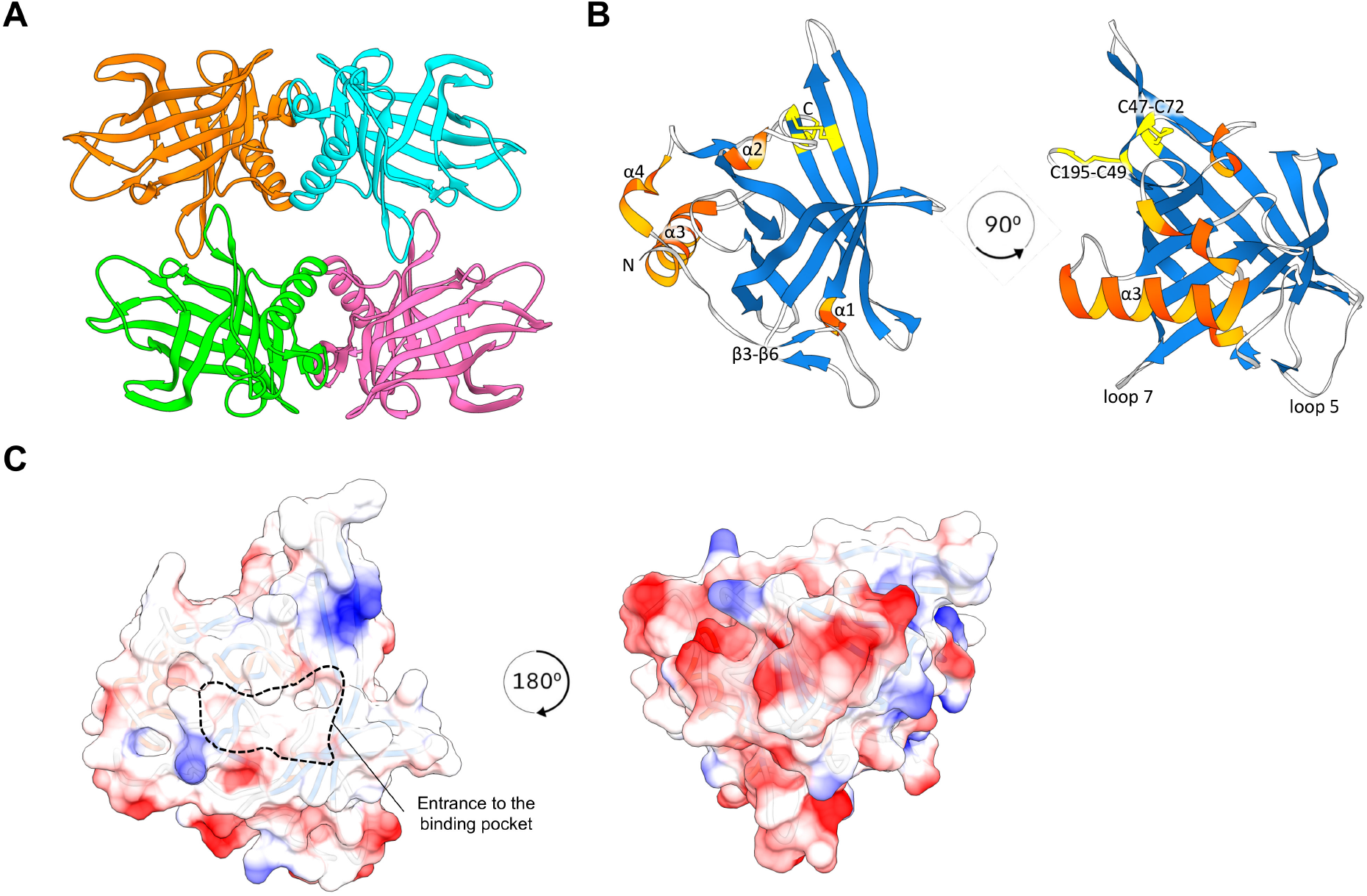
*Pf*LCN forms a tetrameric structure of typical lipocalin monomers. A) Crystal structure of tetrameric *Pf*LCN. B) Crystal structure of monomeric *Pf*LCN. The N- and C-termini are labelled, as are various structural features including the two disulphide bonds. C) Representation of Coulombic surface charges on *Pf*LCN. Red indicates net positive charge regions, blue indicates net negative charge regions. The left view shows the top-down view looking into the binding pocket, while the right view displays a bottom-up view. For details regarding data collection statistics and refinement statistics see also Figure S3.

The overall monomeric structure of *Pf*LCN reveals the classic lipocalin fold with a β-barrel calyx core consisting of eight hydrogen-bonded anti-parallel β-strands and an N-terminal α-helix (α3) perpendicular to the lumen of the barrel (Figure 1B, Movie S2). There is a typical lipocalin binding pocket with a highly hydrophobic environment inside the β-barrel where non-polar ligands usually bind in most lipocalins, for example the many lipophilic ligands of the human lipocalin NGAL (Bao *et al*, 2015). The binding pocket of *Pf*LCN can be measured as approximately 20 Å by 20 Å at the widest span, which is in a similar range to other lipocalins (Campanacci *et al*, 2004; Eichinger *et al*, 2007). The bottom of the barrel is closed by the inter-strand hydrogen bonds between β-strands 3 and 6, and also a 3_10_ helix. This is a feature also shared with other lipocalins. The available protein structure with the highest degree of similarity to *Pf*LCN is the *E. coli* lipocalin Blc (Campanacci *et al*, 2004) with a RMSD of 1.7 Å and a modified version of Blc was used as a molecular replacement model during structure determination. Interestingly, structural comparisons of *Pf*LCN with human lipocalins (Breustedt *et al*, 2006) revealed that amongst these *Pf*LCN has the highest structural similarity with the high-density lipoprotein ApoD (Eichinger *et al*, 2007) (Figure S4) with a RMSD of 1.8 Å. The sequence homology of both Blc and ApoD when compared to *Pf*LCN are nevertheless low, with 16.8% and 19.2% sequence identity respectively.

The surface charges of *Pf*LCN are distributed mostly on the underside of the barrel and are mainly constituted of positively charged residues. However, both the dimerization face and the binding face are largely composed of non-polar residues, allowing both a hydrophobic effect-driven dimerization and also the direction of hydrophobic ligands into the binding pocket, which contains mostly non-polar residues (Figure 1C and D). There are two disulfide bonds located close to each other in each monomer (Figure 1B). One disulfide bond (C49-C195) anchors the C-terminal tail to the body of the protein, which is a feature well conserved among lipocalins (Flower, 1996). However, the other disulfide bond (C47-C72) is more unusual in the family and binds the terminus of α2, the helix that appears to be ‘gating’ the entrance to the binding pocket, to β4, one of the longest strands of the barrel. The binding pocket in the structure is mostly occluded by the α2 helix, and while this does not rule out the possibility of ligand binding in these conditions, the C47-C72 disulfide bond may be contributing to controlling access to the pocket.

To investigate whether the tetramer observed in the crystal structure was a physiologically relevant oligomer, we performed small-angle X-ray scattering (SAXS) experiments. These confirmed the presence of a tetramer in solution, and that this tetramer exhibits a concentration-dependent equilibrium with dimeric *Pf*LCN (Figure 2). The dimer obtained from SAXS modeling spatially superimposes well with the crystallographic dimer, which along with the favorable solvation enthalpy gives an indication that this is a stable, physiological dimer. In the crystallographic tetramer and the SAXS-derived dimer, all binding sites are pointing outwards, allowing each monomer to potentially bind to a ligand in this state and remain functional.

**Figure 2.**
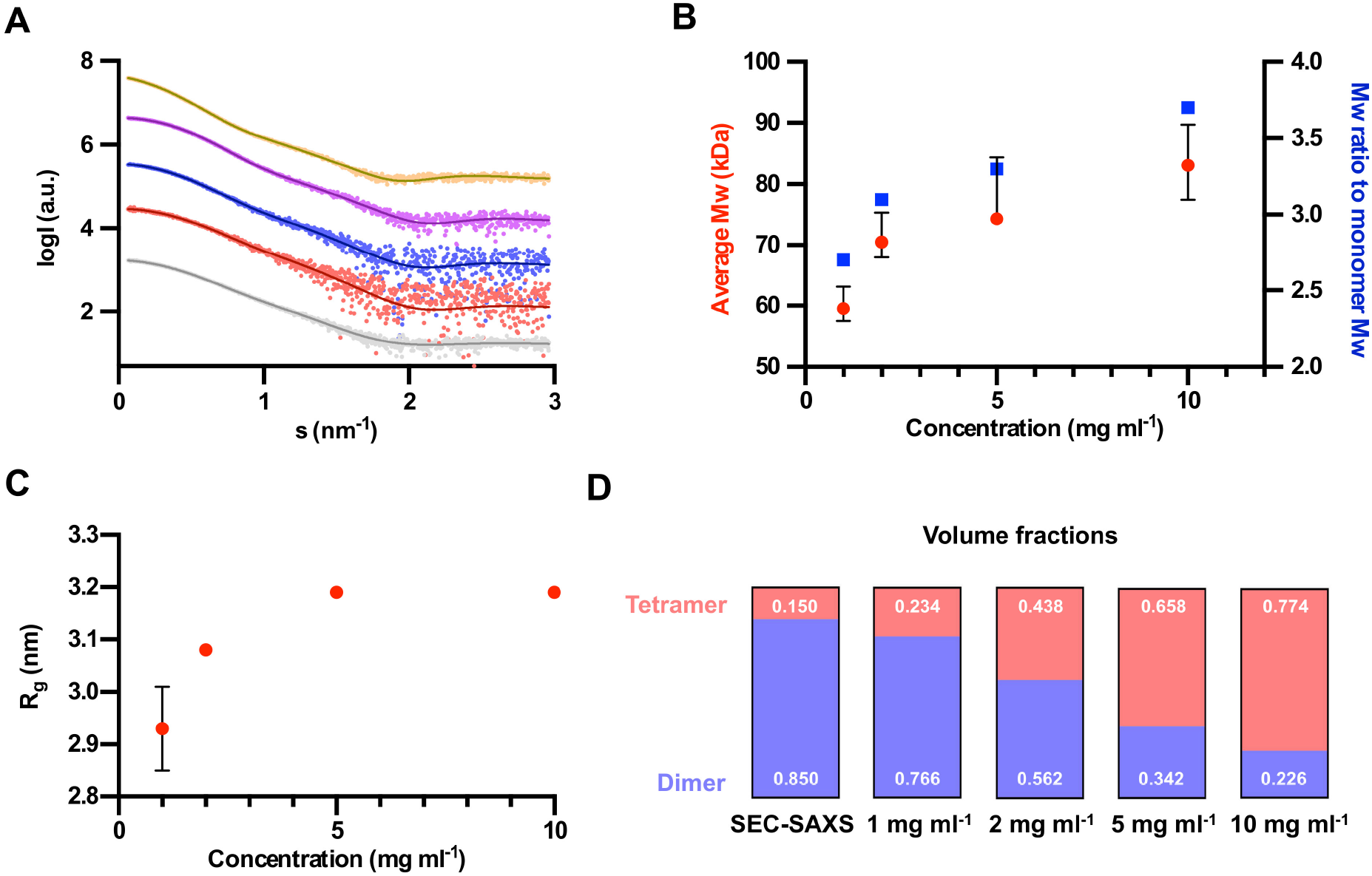
SAXS analysis of the oligomerization of *Pf*LCN in solution. A) SAXS raw data taken over a range of concentration of *Pf*LCN in batch mode, also including a SEC-SAXS run. From bottom to top, the data sets are: SEC-SAXS run, 1 mg ml^−1^, 2 mg ml^−1^, 5 mg ml^−1^ and 10 mg ml^−1^. The fitted lines represent the calculated scattering from the tetramer-dimer mixtures given in Figure 2D, using the software SASREFMX. The Χ^2^ functions of each fit are respectively: 0.98, 0.98, 1.0, 1.2, 1.3. The datasets are offset by a factor of 10 each time for clarity. B) Graph shows concentration of *Pf*LCN plotted against average molecular weight (Mw) of the sample on the left y axis. On the right y axis is the ratio of this Mw compared to the monomer Mw. The SEC-SAXS sample is not plotted on the graph as it is not possible to work out the protein concentration using this technique. C) The average hydrodynamic radius (R_g_) plotted against concentration. D) The proportion of tetrameric and dimeric *Pf*LCN present at different concentrations, calculated using the software SASREFMX.

### *Pf*LCN localizes to the PV and FV and can integrate into membranes

Having shown that *Pf*LCN is indeed a lipocalin based on its structure, we next determined its localization by endogenously tagging *Pf*LCN with green fluorescent protein (GFP) using the selection-linked integration (SLI) system (Birnbaum *et al*, 2017). We additionally introduced a GlmS ribozyme (Prommana *et al*, 2013) sequence upstream of the 3’ untranslated region that allows inducible knockdown of protein expression upon addition of glucosamine to the parasite culture medium (Figure 3A). In parallel, we generated a similar parasite line, in which the GlmS ribozyme sequence was inactivated (M9 mutant (Prommana *et al*, 2013)). Correct integration of both of the corresponding targeting constructs into the endogenous *Pf*LCN locus was confirmed by PCR (Figure 3B). In the following, the resulting parasite lines are referred to as *Pf*LCN-knockdown (LCN-Kd, GlmS-WT sequence) and *Pf*LCN-control parasites (LCN-Con, GlmS-M9 sequence).

**Figure 3.**
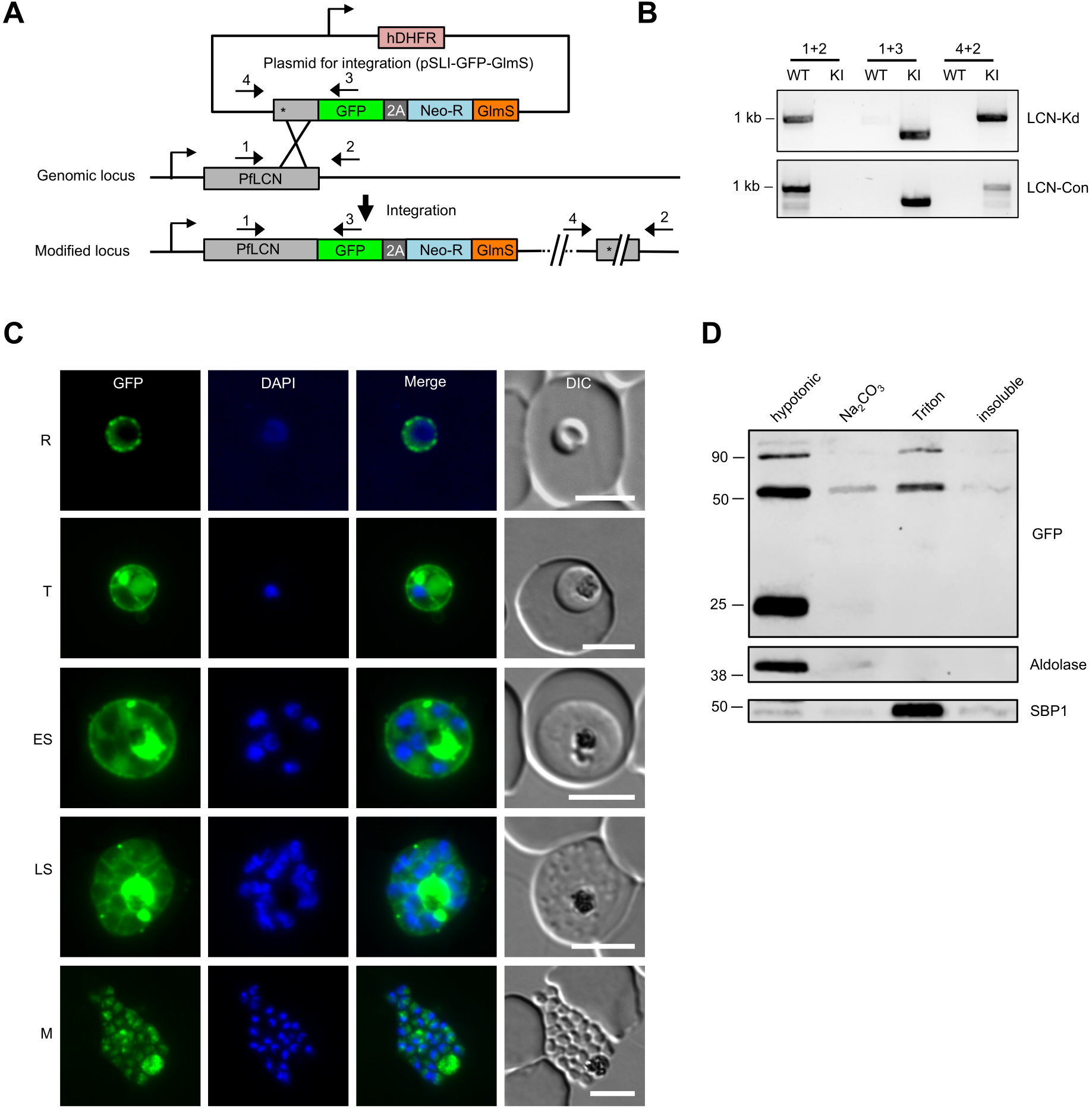
Endogenous tagging and localization analysis of *Pf*LCN. A) Schematic of the selection-linked integration based strategy used to generate LCN-Kd and LCN-Con parasites by single-crossover recombination. Localization of primers used for showing successful integration of targeting constructs by PCR are indicated. T2A, skip peptide; Neo-R, neomycin-resistance gene; GlmS, GlmS ribozyme; asterisks, stop codons; arrows, promoters. B) Integration PCR of transgenic LCN-Kd and LCN-Con knock in (KI) and unmodified wild type (WT) parasites. C) Localization of *Pf*LCN-GFP (green) by live-cell microscopy in ring (R), trophozoite (T), early schizont (ES), late schizont (LS) stage parasites as well as in free merozoite (M). Nuclei were stained with DAPI (blue). DIC = Differential interference contrast. Scale bar = 5 μm.
D) Sequential differential solubility analysis of *Pf*LCN-GFP parasite saponin pellet by hypotonic lysis (soluble proteins), treatment with sodium carbonate (Na_2_CO_3_, peripheral membrane proteins) and Triton X-100 (integral membrane proteins). The soluble protein aldolase and the integral membrane protein skeleton-binding protein 1 (SBP1) served as positive controls. The band at around 25 kD in the soluble fraction likely represents monomeric GFP, the band at around 50 kD is monomeric *Pf*LCN-GFP, while the higher molecular band at around 90 kD is likely derived from dimeric *Pf*LCN-GFP. Solubility results are representative of three independent experiments, of which one is shown.

Live-cell microscopy of LCN-Kd parasites revealed that in rings and early trophozoites *Pf*LCN-GFP localized to the parasite periphery, suggesting a possible localization on the PVM, the PPM or within the PV itself (Figure 3C). In trophozoites and early schizonts, the protein was additionally visible in small, mostly peripheral, focal structures, the FV and in accumulations in the parasite cytoplasm. In schizonts, we mainly observed the *Pf*LCN signal around developing merozoites, in some accumulations in the schizont cytoplasm and in the FV. In contrast, in free merozoites *Pf*LCN was present in the cytosol but no GFP signal was evident around their periphery, excluding a localization to the PPM and further supporting that *Pf*LCN localizes to the PV and possibly on the PVM of the infected erythrocyte (Figure 3C).

Given that several lipocalins, including *E. coli* Blc and human ApoD, have the capability to integrate into membranes (Hernández-Gras & Boronat, 2015; Eichinger *et al*, 2007; Bishop *et al*, 1995), we next investigated the membrane association of *Pf*LCN and performed sequential differential solubilization of lysed parasite membranes. To this aim, saponin (permeabilizes erythrocyte membrane and PVM) pellets of Lip-Kd trophozoite stage parasites were hypotonically lysed to release non-membrane associated proteins and then sequentially treated with sodium carbonate and Triton X-100 to solubilize peripheral and integral membrane proteins, respectively. Western-Blot analysis using anti-GFP antibodies revealed that an approximately 50 kDa *Pf*LCN-GFP (calculated MW 50.8 kDa) was not only found in the soluble fraction but also in the sodium carbonate and Triton fraction, indicating that it is a soluble protein but that it can also associate with and integrate into membranes (Figure 3D). Interestingly, we also detected a higher molecular mass GFP-positive band, which likely represents *Pf*LCN in its dimeric state as previously described for other lipocalins (Bhatia *et al*, 2012; Van Veen *et al*, 2006; Kielkopf *et al*, 2018) and as also supported by our SAXS analysis. In addition to these bands, a third band at around 25 kD was visible that likely represents cleaved GFP. This band is often detected in GFP-fusions in *P. falciparum* in particular for PV and FV proteins (Klemba *et al*, 2004; Boddey *et al*, 2016) and was solely found in the soluble fraction further underlining the specificity of *Pf*LCN’s membrane association.

### *Pf*LCN is important for parasite survival in erythrocytes

In order to test if *Pf*LCN is essential for blood stage proliferation, we first tried to functionally inactivate it by targeted-gene disruption using the SLI system (Birnbaum *et al*, 2017). However, repeated attempts failed to obtain parasites carrying the correctly integrated plasmid, indicating that the gene is likely essential for parasite development.

We next aimed to conditionally downregulate *Pf*LCN expression and to study the influence of this on parasite development. To test whether the GlmS-system can be used to investigate *Pf*LCN function, we first added 2.5 mM glucosamine to tightly synchronized early ring stage LCN-Kd and LCN-Con parasites and measured *Pf*LCN expression 24 hours post glucosamine addition by quantitative western blot analysis. This revealed a >95% knockdown of the protein in LCN-Kd parasites upon glucosamine addition, while no influence on *Pf*LCN expression was seen in LCN-Con parasites, showing efficient regulation of *Pf*LCN expression by the GlmS ribozyme (Figure 4A).

**Figure 4.**
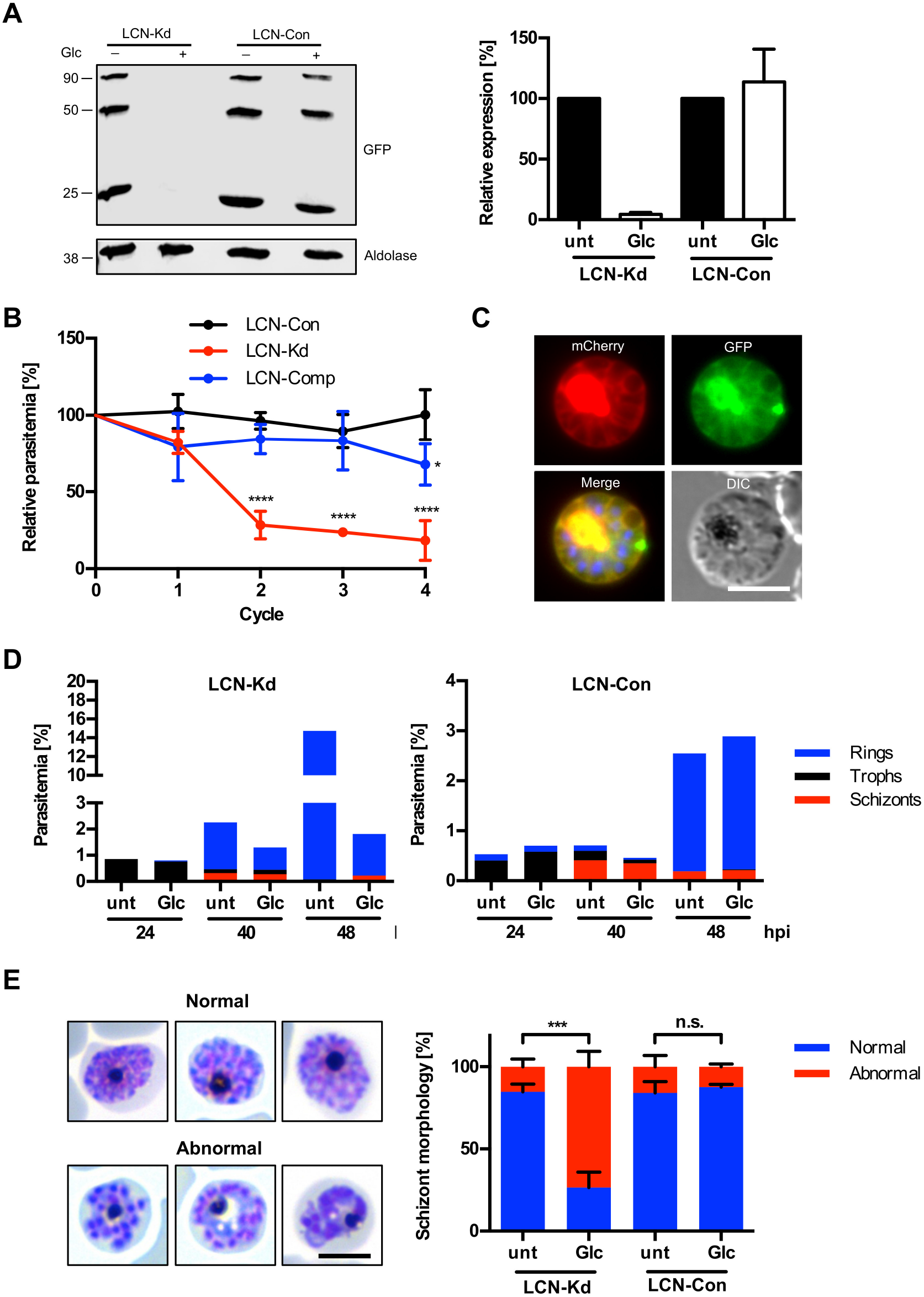
*Pf*LCN is essential for parasite growth and for late stage maturation. A) Western blot of LCN-Kd and LCN-Con trophozoite stage parasites, which had been treated for 24 hours with 2.5 mM glucosamine or that were left untreated. Aldolase served as a loading control. Quantification of monomeric *Pf*LCN-GFP levels normalized to the amount of aldolase is displayed on the right. Shown are means +/− SD of three independent experiments, in which *Pf*LCN-GFP levels of untreated parasites were always set to 100%. The band at around 25 kD most propably represents monomeric GFP, the band at around 50 kD is monomeric *Pf*LCN-GFP, while the higher molecular band at around 90 kD is likely derived from dimeric *Pf*LCN-GFP. B) Growth analysis of LCN-Kd, LCN-Con and complemented LCN-Kd (LCN-Comp) parasites over four parasite cycles in presence or absence of 2.5 mM glucosamine. Shown are relative parasitemia values, which were obtained by dividing the parasitemia of glucosamine treated cultures by the parasitemia of the corresponding untreated ones. Shown are means +/− SD of four independent growth experiments. For statistical evaluation of the growth of LCN-Kd and LCN-Comp parasites in comparison to LCN-Con parasites, a one-way analysis of variance (ANOVA) followed by a Holm-Sidak multiple comparison test was performed. All statistically significant differences are indicated (* p < 0.05, **** p < 0.0001). C) Localization of *Pf*LCN-mCherry in complemented parasites by live-cell microscopy. Nuclei were stained with DAPI (blue). DIC = Differential interference contrast. Scale bar = 5 μm. D) Stage quantifications of LCN-Kd and LCN-Con parasites grown in the presence or absence of 2.5 mM glucosamine at 24, 40 and 48 hpi. Results are representative of three independent experiments, of which one is shown. E) Quantification of schizont morphology of compound 2 arrested LCN-Kd and LCN-Con schizonts grown in the presence or absence of 2.5 mM glucosamine. Shown are means +/− SD of three independent experiments. For statistical evaluation an unpaired two-tailed students t-test was performed. Statistically significant differences are indicated (*** p < 0.001, n.s. = not significant). Representative Giemsa-stained schizonts with normal and abnormal morphology are shown on the left. Scale bar = 5 μm.

To assess the influence of *Pf*LCN knockdown on parasite multiplication, we next quantitated the growth of LCN-Kd and LCN-Con parasites with and without glucosamine using a flow cytometry-based growth assay over four parasite cycles. Addition of glucosamine to LCN-Kd parasites resulted in a pronounced growth defect, which was already visible in the first parasite cycle following glucosamine addition (Figure 4B). In contrast, no noticeable growth defect was evident for LCN-Con parasites treated with glucosamine, thereby excluding that the observed growth defect in LCN-Kd parasites is due to glucosamine cytotoxicity. To further verify that the observed reduction in multiplication of LCN-Kd parasites grown in the presence of glucosamine is due to the knockdown of *Pf*LCN, we complemented LCN-Kd parasites with an episomal plasmid, from which the *Pf*LCN coding sequence is expressed as a fusion with mCherry under the constitutive *nmd3* promoter (Birnbaum *et al*, 2017). As expected, in these parasites the mCherry-tagged *Pf*LCN showed a clear PV localization in addition to a pronounced staining of the parasite FV (Figure 4C). Importantly, when we added glucosamine to these complemented parasites, the growth defect was reversed, confirming that knockdown of *Pf*LCN is specific and deleterious for parasite growth (Figure 4B).

### *Pf*LCN knockdown impairs late stage maturation

To determine which particular parasite stage is affected by the knockdown of *Pf*LCN, we added glucosamine to tightly synchronized LCN-Kd and LCN-Con ring stage parasites and quantified the different parasite stages at 24, 40 and 48 hours post invasion. While we did not see any major effect on trophozoite and schizont numbers upon *Pf*LCN knockdown, there was a substantial reduction of newly formed rings at 48 hpi. As expected, no such effect was seen in LCN-Con parasites after glucosamine addition (Figure 4D). To investigate this further, we analyzed schizont morphology in Giemsa-stained blood smears of untreated and glucosamine treated LCN-Kd and LCN-Con parasites that had been incubated with the egress inhibitor compound 2 from 40 to 48 hpi. Interestingly, there was a statistically significant increase of schizonts with abnormal morphology upon glucosamine addition to LCN-Kd parasites, suggesting that *Pf*LCN knockdown affects late stage maturation of the parasite (Figure 4E).

### *Pf*LCN function affects hemozoin crystal motility in the FV and is related to the activity of ROS

To understand the late stage maturation defect of *Pf*LCN knockdown parasites in more detail, we also studied LCN-Kd and LCN-Con trophozoite stage parasites by live-cell microscopy. Interestingly, glucosamine addition to LCN-Kd but not to LCN-Con parasites nearly completely abolished the motion of hemozoin crystals that under normal circumstances move within the food vacuole of individual parasites (Figure 5A, Movies S3 and S4). Hemozoin dynamics serve as internal biomarker for food vacuole integrity and parasite viability, although the origin and physiological significance of this motion is unknown so far (Sigala & Goldberg, 2014). A loss of hemozoin crystal motility was previously observed by photoillumination of the heme synthesis intermediate protoporphyrin X due to the generation of ROS (Sigala *et al*, 2015). Based on this, we hypothesized that the observed loss of hemozoin crystal motility in *Pf*LCN knockdown parasites is connected to ROS activity. To test this, we induced the knockdown of *Pf*LCN in LCN-Kd parasites in the presence of the radical scavenger Trolox, a water-soluble analog of vitamin E. Remarkably, this resulted in a partial rescue of the hemozoin crystal motility phenotype, implying ROS mediated oxidation in the loss of hemozoin motility (Figure 5A). In line with this, treatment of LCN-Kd parasites with the thiol-reactive chemical diamide, which has been shown to induce oxidative stress in the parasite (Rahbari *et al*, 2017), also inhibited hemozoin crystal motility. Again, this drug induced motility arrest could partially be restored by the addition of Trolox (Figure 5B).

**Figure 5.**
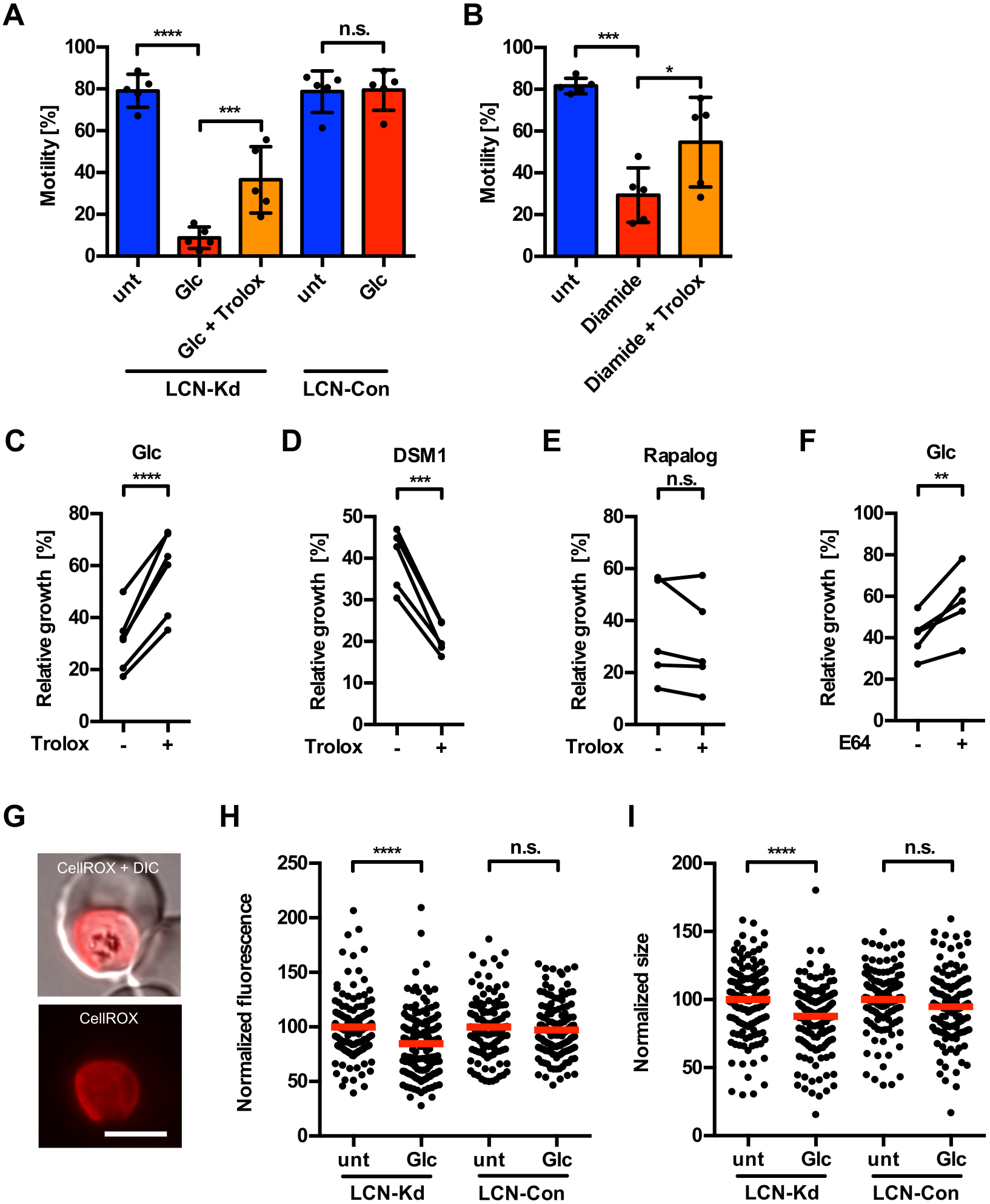
*Pf*LCN protects parasites against oxidative stress induced cell damage. A) Hemozoin crystal motility analysis of LCN-Kd and LCN-Con trophozoite stage parasites grown in the presence or absence of 2.5 mM glucosamine. In a separate culture, 100 uM of the radical scavenger Trolox was added to LCN-Kd knockdown parasites. B) Hemozoin crystal motility analysis of LCN-Kd trophozoite stage parasites to which 10 μM diamide in presence or absence of 100 μM Trolox was added for 4 hours before analysis. Shown are means +/− SD of five independent experiments. For statistical comparisons a one-way ANOVA followed by a Holm-Sidak multiple comparison test was performed. Statistically significant differences are indicated (* p < 0.05, *** p < 0.001, **** p < 0.0001, n.s. = not significant). Please see also Movies S3 and S4. C, D) Growth analysis of LCN-Kd parasites over one parasite cycle grown in presence or absence of 2.5 mM glucosamine or 0.24 μM DSM1 to which 100 μM Trolox were added or that were left untreated. E) Growth analysis over one parasite cycle of Rab5a knocksideways parasites grown in presence or absence of 40 nM Rapalog to which 100 μM Trolox were added or that were left untreated. Note that the amounts of DSM1 and Rapalog were chosen to mimic approximately the reduction in growth observed upon *Pf*LCN knockdown. F) Growth analysis of LCN-Kd parasites over one parasite cycle grown in presence or absence of 2.5 mM glucosamine to which 1 μM E64 was added for 12 hours at the ring stage or that were left untreated. In C,D,E,F relative parasitemia values of five to six independent experiments are displayed, which were obtained by dividing the parasitemia of glucosamine/DSM1/Rapalog treated cultures by the parasitemia of the corresponding untreated ones both in presence or absence of Trolox/E64. For statistical analysis of these experiments a ratio-paired two-tailed t-test was performed. Statistically significant differences are indicated (** p < 0.01, *** p < 0.001, **** p < 0.0001, n.s. = not significant). G) Live-cell image of an LCN-Kd parasite incubated with the ROS-sensitive dye CellROX orange. DIC = Differential interference contrast. Scale bar = 5 μm. H) Oxidative stress levels in LCN-Kd and LCN-Con trophozoite stage parasites grown in presence or absence of 2.5 mM gluosamine as measured by CellROX Orange, which shows enhanced red fluorescence upon higher levels of ROS. Parasite sizes (area) were measured in parallel and are displayed in (I). Shown are pooled data from three independent experiments in which a total of 114-151 parasites were analyzed per cell line and condition. Mean values are highlighted in red. In each experiment, mean fluorescence levels and parasite sizes were set to 100% in untreated LCN-Kd and LCN-Con parasites. For statistical analysis an unpaired two-tailed students t-test was performed. Statistically significant differences are indicated (**** p < 0.0001, n.s. = not significant).

### *Pf*LCN reduces oxidative cell damage

To further analyze the importance of ROS, we induced the knockdown of *Pf*LCN in the presence or absence of Trolox. Concordantly with the rescue of hemozoin crystal motility, Trolox treatment chemically complemented around 50% of the growth defect of *Pf*LCN knockdown parasites (Figure 5C), suggesting that *Pf*LCN function might be associated with protection from ROS or ROS-induced cell damage. In order to control for the possibility that Trolox treatment non-specifically protects parasites from cell death, we also treated them with the drug DSM1, which targets dihydroorotate dehydrogenase in the mitochondrial electron transport chain leading to inhibition of pyrimidine biosynthesis (Phillips *et al*, 2008). However, Trolox treatment failed to restore growth of parasites treated with DSM1 and instead potentiated the activity of the drug (Figure 5D). As another control, we conditionally mislocalized the GTPase Rab5a since inactivation of this protein causes a similar phenotype to *Pf*LCN with a defect in late stage parasite maturation (Birnbaum *et al*, 2017). Importantly and in contrast to *Pf*LCN knockdown, no difference in growth was observed upon Trolox treatment here (Figure 5E), further highlighting the specificity of the Trolox-based rescue of parasite growth upon *Pf*LCN knockdown.

One main source of ROS in infected erythrocytes is the degradation of hemoglobin within the parasite FV (Atamna & Ginsburg, 1993) and this process can be inhibited by application of protease inhibitors (Atamna & Ginsburg, 1993; Mesén-Ramírez *et al*, 2019). We therefore tested whether temporal inhibition of hemoglobin digestion through the addition of the cysteine protease inhibitor E64 similarly rescues growth of *Pf*LCN knockdown parasites. To do so, 1 μM E64 was added to ring stage LCN-Kd parasites in presence or absence of glucosamine with E64 being washed away after 12 hours, a treatment that was previously shown to be effective in inhibiting hemoglobin digestion (Mesén-Ramírez *et al*, 2019). Temporal E64 application also led to a statistically significant rescue of the *Pf*LCN knockdown-induced growth inhibition (Figure 5F), which combined with the ability of Trolox to rescue growth of *Pf*LCN knockdown parasites supports that this protein might have a role in protection from ROS that is produced during hemoglobin digestion in the FV.

Protection from ROS could occur through direct lowering of ROS levels or by mitigating ROS induced cell damage. To differentiate these possibilities, we measured ROS levels in untreated and glucosamine treated LCN-Kd and LCN-Con trophozoite stage parasites by determining the fluorescence intensity of the ROS-sensitive dye CellROX Orange (Figure 5G). Interestingly, ROS levels were slightly lower in glucosamine treated LCN-Kd parasites in comparison to untreated parasites, while no such effect was visible in glucosamine treated LCN-Con parasites (Figure 5H). In parallel to measuring ROS levels, parasite size was also measured in *Pf*LCN knockdown parasites showing a slight decrease in size (Figure 5I). It is possible that this difference in size occurs as a result of impaired food vacuole physiology, which may influence CellROX staining intensity as well. Nevertheless, the fact that *Pf*LCN knockdown did not lead to an increase in parasite ROS compared to wildtype parasites during trophozoite development, suggests that it most likely has a role in reducing oxidative stress induced cell damage.

## DISCUSSION

Lipocalins are a large family of small proteins that play important roles in various different physiological functions. They are part of the larger calycin protein superfamily and share a highly conserved three-dimensional structure but show unusually low levels of sequence similarity. Due to the extreme sequence diversity amongst lipocalins with pairwise identities falling below 20%, it remains challenging to reliably predict new members of the lipocalin family in genomes using sequence search methods. Predictions that rely on a single computational tool may even miss some of the well-characterized lipocalin family members and therefore, they need to be complemented by different methods. Machine learning approaches have been proposed (Ramana & Gupta, 2009), but the currently available algorithms proved unreliable in this study and failed to identify the *Pf*LCN sequence as a lipocalin (Ramana & Gupta, 2009). Here, using a targeted search informed by libraries of protein signatures and hidden Markov models that are implemented in integrated database resources, we identified a new essential member of the lipocalin protein family in *P. falciparum* malaria parasites. We confirmed that *Pf*LCN has a typical lipocalin fold through crystallization and structure solution of the protein, which revealed it to be a novel member of the lipocalin family and the first lipocalin characterized in a single-celled eukaryote.

Remarkably, when the crystal structure of *Pf*LCN was solved it was found to be present as a tetramer containing a dimer of dimers. Using SAXS analysis, we confirmed that *Pf*LCN also forms a tetramer in solution and is in a concentration-dependent equilibrium with its dimeric form. This observation excludes that tetramerization is an artefact of the crystallization process. Oligomerization is a common feature of lipocalins and often occurs upon ligand binding (Kielkopf *et al*, 2018). The fact that *Pf*LCN was in an oligomeric state in the crystal structure and in solution without a ligand bound suggests that the multimerization of *Pf*LCN occurs independently of its ligand. Interestingly, a higher molecular mass GFP-positive band was also observed in addition to the one corresponding to the monomeric protein, when parasites expressing *Pf*LCN-GFP were analysed by western blotting using GFP antibodies. This band is most likely derived from dimeric *Pf*LCN-GFP and points to a high stability of the putative *Pf*LCN-GFP dimer, as it was not destroyed by the reducing conditions of the SDS-PAGE. In addition, it may indicate that *Pf*LCN can also dimerize *in vivo*. In this respect it is interesting to note that reducing SDS-PAGE resistant dimers have already been observed for other lipocalins (Bhatia *et al*, 2012; Van Veen *et al*, 2006; Kielkopf *et al*, 2018).

*Pf*LCN localizes to the main membrane bound vacuolar compartments of the parasite, the PV and the FV. Both compartments are highly sophisticated and have specialized roles that are essential for blood-stage parasite growth. The PV is a bubble-like compartment that is initiated during the process of invasion through the invagination of the erythrocyte membrane. It is bounded by the PVM, which constitutes the parasite-host cell interface and undergoes massive re-modelling after erythrocyte infection is established. The PVM plays a central role in nutrient acquisition, host cell remodelling, waste disposal, environmental sensing, and protection from innate defence mechanisms (reviewed in (Spielmann *et al*, 2012)). Likewise, the FV is a unique compartment that is optimized for hemoglobin metabolism. It is the site of acidification, hemoglobin proteolysis, peptide transport, heme polymerization and detoxification of oxygen radicals (reviewed in (Wunderlich *et al*, 2012)). In contrast to other organelles (e.g. mitochondrion, apicoplast) the FV, like the PV, does not persist throughout the whole asexual intra-erythrocytic life cycle. It is formed *de novo* and is discarded at the end of each cycle.

The PV and FV localization of *Pf*LCN are in agreement with a recent study, in which the *Pf*LCN orthologue in the rodent malaria model *P. berghei* (termed PV5) was identified as an essential PV protein (Matz & Matuschewski, 2018). The FV localization is further supported by mass spectrometric analysis, which showed that *Pf*LCN is part of the FV proteome (Lamarque *et al*, 2008). Given these localizations and the well-established function of lipocalins to transport small hydrophobic molecules (Flower, 1996), *Pf*LCN may well have a role in transport of hydrophobic molecules (e.g. lipids, fatty acids) from or to the PV/FV. Supporting this, *Pf*LCN has a high structural similarity with the bacterial lipocalin Blc, an *E. coli* outer membrane lipoprotein, which has been shown to bind fatty acids and lyso-phospholipids and is most likely involved in the transport of these in the bacterium (Campanacci *et al*, 2006). However, confirmation of a role for *Pf*LCN in small hydrophobic molecule transport, particularly around the PV, will be addressed in future studies as our results suggest that this protein has another essential role centred on the FV.

Our finding that *Pf*LCN localises to the FV puts in context some of our central observations of the protein knockdown: the loss of hemozoin crystal motility within the FV and the rescue of this motility with a radical scavenger. When we compared the structure of *Pf*LCN to human members of the lipocalin family, it became evident that *Pf*LCN shows high structural similarity to human ApoD. Interestingly, the tetramerization of ApoD under physiological condition has recently been described (Kielkopf *et al*, 2018). Human ApoD is capable of binding to a range of small hydrophobic ligands such as progesterone and arachidonic acid and exerts an antioxidant function, which is partially due to the reduction of peroxidized lipids (Bhatia *et al*, 2012). This antioxidative function is conserved in ApoD homologs in mice, flies and in the plant *Arabidopsis thaliana* (Ganfornina *et al*, 2008; Sanchez *et al*, 2006; Walker *et al*, 2006; Charron *et al*, 2008). Supporting a similar antioxidative function of *Pf*LCN at the FV, the *Pf*LCN knockdown induced proliferation phenotype could be partially chemically complemented with the radical scavenger Trolox and by temporal inhibition of hemoglobin digestion using the cysteine protease inhibitor E64. ROS levels in *Pf*LCN knockdown parasites were, however, slightly lower in comparison to control parasites. This might be related to the observation that *Pf*LCN knockdown arrests the motion of hemozoin crystals that, under normal circumstances, move dynamically within the FV of individual parasites. This in turn might be a sign of disordered FV physiology, likely associated with reduced hemoglobin digestion and lower amounts of ROS being produced. In addition to this, ROS levels might be influenced by a *Pf*LCN-knockdown associated effect on parasite growth during trophozoite development, as parasite sizes that were measured in parallel to ROS determination, were also slightly reduced in *Pf*LCN knockdown parasites.

Importantly, the lower ROS levels we observed in knockdown parasites argue against a direct function of *Pf*LCN in scavenging of ROS and would instead favour a function similar to human ApoD in mitigating oxidative stress associated cell damage. Human ApoD catalyzes the reduction of peroxidized lipids via a highly conserved methionine residue. Specifically, it reduces peroxidized, potentially radical-generating hydroperoxy eicosatetraenoic acids to the non-reactive hydroxide forms, whereby lipid peroxidation chain reactions are prevented (Bhatia *et al*, 2012; Dassati *et al*, 2014). *Pf*LCN also contains three relatively conserved methionine residues, two of which are within its binding pocket (Figures S1 and S4). Therefore, we can speculate that a similar mechanism as described for ApoD might also be used by *Pf*LCN to limit oxidative damage, particularly in the FV. This potential membrane protective effect agrees with our observation that *Pf*LCN not only exists as a soluble protein, but that it also associates with and integrates into membranes. Other lipocalins such as human ApoD attach to the membrane by forming a disulphide bond with a membrane-embedded protein, but also use a hydrophobic loop at the opening of the binding pocket to anchor themselves into a membrane (Eichinger *et al*, 2007; Hernández-Gras & Boronat, 2015). *Pf*LCN also contains a highly hydrophobic loop (residues 76-79 between β4-β5) that could be used in the same way. This loop contains two residues of the aromatic amino acid phenylalanine, which is often a residue found inserted into membranes (Wang *et al*, 2009). The apparent ability of *Pf*LCN to protect against oxidative damage might explain why *Pf*LCN knockdown parasites show an aberrant late stage development, when massive amounts of new membrane material has to be synthesized and protected from oxidative damage that is the result of Hb digestion in the FV. How this protection may occur on a molecular level and how it is related to the localization of the protein requires further investigation.

Although we have shown in our experiments using the radical scavenger Trolox that one important function of *Pf*LCN is likely its protection against oxidative stress induced cell damage, we only observed a 50% rescue of the *Pf*LCN deletion phenotype upon Trolox treatment. This might on the one hand be due to the fact that not all ROS are scavenged by Trolox at the same efficiency (Sueishi *et al*, 2014). On the other hand, it might well be possible that *Pf*LCN has other independent functions from its antioxidative effects, such as fatty acid or lipid transport, which contribute to the formation of abnormal schizonts observed in this study.

Here, we have characterized the first lipocalin in protists that we identified in the Apicomplexan malaria parasite *P. falciparum*, further broadening the organisms known to possess members of this protein family. Lipocalins have been hypothesized to share an early evolutionary origin in the prokaryotic world, followed by an evolutionary history that has been shaped by gene duplication, gene loss and extreme amino acid divergence in particular lineages (Ganfornina *et al*, 2000). Since only a few lipocalin genes have been identified in unicellular eukaryotes and such lipocalins do not align well with the ones found in higher eukaryotes, previous studies provide only limited information on putative evolutionary trajectories. Likewise, our attempts to reconstruct gene phylogenies from highly divergent primary sequences did not provide useful evolutionary insights. We therefore aimed to assess the representation of lipocalins across the tree of life from available whole genome datasets, focussing on extant alveolate taxa. Interestingly, when applying the same search strategy used to identify *Pf*LCN in order to identify putative lipocalins in other alveolate species, we found that many parasitic species either lost or encode only one to few putative lipocalin genes, while in most of their free-living photosynthetic relatives a higher number of putative lipocalin genes can be found, taking different genome sizes into account. Although we cannot rule out intra-species duplications, this finding is in agreement with the paradigm that the transition from free-living to parasitic lifestyle is predominantly characterized by gene loss that has very likely also affected the lipocalin family. Whether or not the lipocalins of other Apicomplexans also play a role in reducing oxidative damage, potentially as a result of host-immune responses (Sorci & Faivre, 2009), remains to be determined.

In conclusion, using targeted sequence analysis and by solving its structure using X-ray crystallography, we were able to confirm the identity of the first lipocalin in a protist. We show that *Pf*LCN is an essential protein of the malaria parasite *P. falciparum*, which localizes to the PV and FV. We functionally link *Pf*LCN to oxidative damage control and provide evidence that the antioxidative function of lipocalins not only occurs in mammals, insects and plants but also in a unicellular eukaryote.

## MATERIALS AND METHODS

### Identification and evolutionary analysis of lipocalin superfamily in *Plasmodium falciparum* and related protists

Searching the PlasmoDB database (https://plasmodb.org/) for genes annotated as members of the lipocalin superfamily (SSF50814; SUPERFAMILY database (Pandurangan *et al*, 2019)) or the calycin homologous superfamily (IPR012674; InterPro (Mitchell *et al*, 2019)) revealed the protein coding gene PF3D7_0925900 as putative new lipocalin. We performed batch blastp searches of *Pf*LCN (PF3D7_0925900) against the nr and env_nr database (Agarwala *et al*, 2018) using Geneious 10.2.3 (https://www.geneious.com) using an E-value of 10e-0 (BLOSUM62 substitution matrix) to identify close homologs in related apicomplexan species. A multiple sequence alignment of putative lipocalins identified in *Plasmodium* species was performed using MUSCLE implemented in the R package msa v1.16.0 (Bodenhofer *et al*, 2015).

We then used the InterPro (Mitchell *et al*, 2019) and SUPERFAMILY 2.0 database (Pandurangan *et al*, 2019) to identify and count the number of lipocalin superfamily members in selected alveolate and outgroup species to perform an evolutionary analysis. The R package rotl v3.0.10 (Michonneau *et al*, 2016) was used to extract a phylogenetic tree from the Open Tree of Life v11.4 synthetic tree (Hinchliff *et al*, 2015) summarizing current phylogenetic and taxonomic information. Additionally, we obtained genome size data from the NCBI genome database (Agarwala *et al*, 2018) (https://www.ncbi.nlm.nih.gov/genome/browse) and visualized the phylogenetic tree and associated protein count and genome size data using the R package ggtree v1.16.6 (Yu *et al*, 2018).

### Protein expression and purification

The coding sequence of *Pf*LCN (corresponding to residues 22-217 and lacking the 21 amino acid N-terminal signal sequence) was PCR amplified from blood stage cDNA using primers rec*Pf*LCN-fw and rec*Pf*LCN-rev and cloned into the pCoofy1 vector (Scholz *et al*, 2013) containing an N-terminal hexahistidine tag followed by a 3C PreScission protease site (sequence: LEVLFQG) using the SLiCE cloning method (Zhang *et al*, 2014). The vector was transformed into Rosetta-gami 2 (DE3) cells using standard protocols. Cells were grown to an OD_600_ of 1.0 in Terrific Broth media at 37°C before induction with 0.5 mM IPTG at 20°C for 18 hours. Cells were pelleted by centrifugation (9373 g) and resuspended in lysis buffer (20 mM Tris pH 7.5, 100 mM NaCl, 20mM imidazole, 5% (v/v) glycerol) before lysis with the Avestin EmulsiFlex C3 french press. Unlysed cells and precipitates were removed by centrifugation (29616 g) and the supernatant was collected and filtered. Filtered lysate was then flowed over Ni-NTA agarose resin at 2.5 ml min^−1^ before a 7-column volume wash with lysis buffer. *Pf*LCN was eluted using a step gradient with elution buffer (20 mM Tris pH 7.5, 100 mM NaCl, 500 mM imidazole, 5% (v/v) glycerol). Fractions were collected and pooled and 1 mg 3C protease was added. The *Pf*LCN and 3C protease mixture was dialysed against 20 mM Tris pH 7.5, 100 mM NaCl, 5% (v/v) glycerol using 8 kDa MW cut-off dialysis tubing overnight at 4°C. Reverse Ni-NTA was performed to remove cleaved hexahistidine tag and 3C protease and the flowthrough was concentrated in a 10 kDa Amicon Ultra centrifugal filter. Protein was injected onto a HiLoad 16/60 Superdex 75 prep grade size exclusion chromatography column running dialysis buffer at 1 ml min^−1^. Fractions were pooled and concentrated using the same Amicon concentrator. The protein concentration was measured using OD_280_ on a Nanodrop using the extinction coefficient calculated from the primary sequence before flash freezing and storage at −80°C. All primer sequences are listed in Table S1.

### Crystallization and data collection

Crystallization screenings were carried out using the sitting-drop vapor diffusion method in Swissci plates and with a Mosquito nanolitre-dispensing crystallization robot. 25 mg/ml of recombinant *Pf*LCN in precipitant to protein volume ratios 1:2, 1:1, 2:1 were used for screening in QIAGEN JCSG Core I-IV screens. The crystals appeared around 30 days in 0.1 M Tris pH 7, 0.2 M MgCl_2_, 2.5 M NaCl, and these were cryoprotected in the same buffer plus 20% glycerol. X-ray diffraction data were collected at 100° K on beamline P13 (PETRA III, EMBL Hamburg) using a wavelength of 0.976 Å.

### Structure determination

All work was performed using the CCP4i2 (Potterton *et al*, 2018) suite unless stated. The dataset was integrated using XDS (Kabsch, 2010) and scaled using AIMLESS (Evans & Murshudov, 2013). The solution of the tetrameric asymmetric unit was obtained by molecular replacement using Phaser (McCoy *et al*, 2007) with a model generated and modified by the Phenix MRage pipeline (Bunkóczi *et al*, 2013) based on Blc from *E. coli* (PDB code 3MBT). The initial model was built by Buccaneer (Cowtan, 2006), refined in real space with Coot (Emsley *et al*, 2010) and reciprocal space with REFMAC5 (Murshudov *et al*, 2011). Structure was validated using Molprobity (Williams *et al*, 2018) and displayed using Chimera (Pettersen *et al*, 2004).

### SAXS and SEC-SAXS

Synchrotron SAXS data were measured from the *Pf*LCN protein through a concentration series using a standard batch setup as well as in-line size-exclusion chromatography SAXS (SEC-SAXS). All samples were reconstituted in 20 mM Tris pH 7.5, 100 mM NaCl, 5% v/v glycerol. The data were collected on the EMBL P12 beamline at PETRA III (Hamburg, Germany) equipped with Pilatus 6M detector at a sample-detector distance of 3 m and at a wavelength of λ = 0.124 nm (I(s) vs s, where s = 4πsinθ/λ, and 2θ is the scattering angle). For the batch measurements, the sample concentrations were: 10, 5, 2 and 1 mg/ml (20 successive 0.050 second frames were collected for each sample at 10° C, in addition to the corresponding matched solvent blank). The data were normalized to the intensity of the transmitted beam and radially averaged and the scattering of the solvent-blank was subtracted to generate the SAXS data displayed in this entry. The SEC parameters were as follows: A 75 μl sample at 10 mg/ml was injected at a flow rate of 0.35 ml/min onto a GE Superdex 200 Increase 10/300 column at 20°C. 56 successive 1 second frames were collected through the major SEC-elution peak and processed using CHROMIXS that included the subtraction of an appropriate solvent blank measured from the protein-free column eluate.

### *P. falciparum* culture and transfection

Blood stages of *P. falciparum* parasites (strain 3D7) were cultured in human RBCs (O+) (transfusion blood, Universitätsklinikum Hamburg Eppendorf, Hamburg). Cultures were maintained at 37° C in an atmosphere of 1% O_2_, 5% CO_2_, and 94% N_2_ and cultured using RPMI complete medium containing 0.5% Albumax according to standard procedures (Trager & Jensen, 1976). For transfection of constructs, 60% Percoll-enriched synchronized mature schizonts were electroporated with 50 μg of plasmid DNA using a Lonza Nucleofector II device (Moon *et al*, 2013). Selection was done either with 3 nM WR99210 or 0.9 μM DSM1 (BEI Resources; https://www.beiresources.org). For generation of stable integrant cell lines, parasites containing the episomal plasmids selected with WR were grown with 400 μg/ml G418 to select for integrants carrying the desired genomic modification as described previously (Birnbaum *et al*, 2017).

### Cloning of constructs for parasite transfection

To generate the plasmid pSLI-*Pf*LCN-TGD, 258 bp of the n-terminal *Pf*LCN coding sequence was amplified by PCR from genomic DNA using primers *Pf*LCN-TGD-fw and *Pf*LCN-TGD-rev and cloned into pSLI-TGD-GFP (Birnbaum *et al*, 2017) using NotI and MluI restriction sites.

To generate pSLI-GFP-GlmS-WT and pSLI-GFP-GlmS-M9, the glmS-WT and glmS-M9 sequences were amplified by PCR from pARL-HA-glmS (Flammersfeld *et al*, 2019) using primers GlmS-WT-fw/GlmS-rev (WT) or GlmS-M9-fw/GlmS-rev (M9) and cloned via XhoI into pSLI-TGD-GFP (Birnbaum *et al*, 2017). Next, 498 bp of the c-terminal *Pf*LCN coding sequence omitting the stop codon were amplified by PCR from genomic DNA using *Pf*LCN-GFP-fw and *Pf*LCN-GFP-rev and cloned via NotI and MluI into pSLI-GFP-GlmS-WT and pSLI-GFP-GlmS-M9 to generate the final targeting constructs pSLI-*Pf*LCN-GFP-GlmS-WT and pSLI-*Pf*LCN-GFP-GlmS-M9.

For generation of the *Pf*LCN gene complementation vector pNMD3:*Pf*LCN-mCherry-DHODH, the *Pf*LCN coding sequence without stop codon was amplified by PCR from genomic DNA using primers *Pf*LCN-Comp-fw and *Pf*LCN-Comp-rev and cloned via XhoI and SpeI into the pNMD3:1xNLS-FRB-mCherry-DHODH plasmid (Birnbaum *et al*, 2017) replacing the 1xNLS-FRB sequence with the *Pf*LCN coding sequence. All primer sequences are listed in Table S1.

### Generation of Lip-Kd and Lip-Con parasites and *Pf*LCN gene complementation

The two targeting constructs pSLI-*Pf*LCN-GFP-GlmS-WT and pSLI-*Pf*LCN-GFP-GlmS-M9 were transfected as described above into *P. falciparum* 3D7 parasites, which were then subjected to sequential WR and G418 selection resulting in LCN-Kd (GlmS-WT) and LCN-Con (GlmS-M9) parasites. To confirm correct integration, genomic DNA from parasites selected under G418 as well as from 3D7/WT parasites was prepared with the QIAamp DNA Mini Kit and analyzed by PCR using primers specific for the 5’ and 3’ integration junctions of the *Pf*LCN locus and primers to detect the original locus. For gene complementation of *Pf*LCN, LCN-Kd parasites were transfected with the pNMD3:*Pf*LCN-mCherry-DHODH plasmid and transgenic LCN-Comp parasites were selected by DSM1. All primer sequences are listed in Table S1.

### Live cell microscopy

For live cell microscopy, LCN-Kd parasites were incubated with 1 μg/ml DAPI in culture medium for 15 minutes at 37°C to stain nuclei. Parasites were then washed once in PBS and imaged in PBS on a Leica D6B fluorescence microscope equipped with a Leica DFC9000 GT camera and a Leica Plan Apochromat 100x/1.4 oil objective. Image processing was performed using ImageJ.

### Solubility analysis of *Pf*LCN

For determination of membrane association, trophozoite stage LCN-Kd parasites from two 10 ml dishes (4% hematocrit, 5-10% parasitemia) were isolated using a Percoll gradient as previously described (Heiber & Spielmann, 2014) and released from RBCs using 0.03% saponin in 200 μl of PBS for 10 minutes (if not otherwise indicated, all steps were carried out on ice). After centrifugation at 16.000 g for 10 minutes, the pellet was 3x washed in PBS, complete protease inhibitor (Roche) was added to the pellet and the parasites were lysed in 200 μl of 5 mM Tris-HCl pH 8.0 and frozen at −80°C. After another three freeze thaw cycles in liquid nitrogen, extraction was performed sequentially: The lysate was centrifuged at 16.000 g for 10 min. The supernatant was removed, recentrifuged for 5 min to remove residual insoluble material and saved as the soluble protein fraction. The pellet of the hypotonic lysis was resuspended in 200 μl of freshly prepared 0.1 M Na_2_CO_3_ and stored on ice for 30 min to extract peripheral membrane proteins. After centrifugation, the supernatant was saved, the pellet extracted for 30 min with 200 μl 1% Triton X-100 and centrifuged to obtain the integral membrane protein fraction in the supernatant. The pellet was resuspended in PBS, representing the insoluble fraction. Reducing SDS sample buffer was added to all supernatants as well as to the insoluble fraction and equal amounts of all samples were analyzed by SDS-polyacrylamide gel electrophoresis (PAGE) and western blotting.

### Immunoblotting analysis

Protein samples were resolved by SDS-PAGE and transferred to nitrocellulose membranes (LICOR). Membranes were blocked in 5% milk in TBS-T followed by incubation in the following primary antibodies that were diluted in TBS-T containing 5% milk: mouse-anti-GFP (Sigma, 1:1000), rabbit-anti-aldolase (Mesén-Ramírez *et al*, 2016) (1:2000), mouse-anti-SBP1-N (Mesén-Ramírez *et al*, 2016) (1:2000). After 3x washing in TBST-T, membranes were incubated in similarly diluted secondary antibodies: goat-anti-mouse-800CW (LICOR, 1:10.000), goat-anti-rabbit-680RD (LICOR, 1:10.000) and goat-anti-mouse-680RD (LICOR, 1:10.000). Subsequently, membranes were washed another 3 times with TBST-T and scanned on a LICOR Odyssey FC imager.

### *Pf*LCN knockdown and phenotypic characterization

To induce knockdown of *Pf*LCN, 2.5 mM glucosamine was added to highly synchronous ealy rings stage LCN-Kd parasites. As a control, the same amount of glucosamine was also added to LCN-Con parasites. For all analyses, medium was changed daily and fresh glucosamine and/or compounds were added every day.

For quantification of knockdown efficiency, untreated and gluocosamine treated LCN-Kd and LCN-Con trophozoite stage parasites from one 10 ml dish (4% hematocrit, 5-10% parasitemia) were isolated using a Percoll gradient 24 hours after glucosamine addition as previously described (Heiber & Spielmann, 2014), directly lysed in reducing SDS-sample buffer and analyzed by SDS-Page and western blotting as explained above. Quantifications of *Pf*LCN-GFP and aldolase levels were performed using the LICOR Image Studio software.

For growth analyses over four parasite cycles, highly synchronous early ring stage parasites were diluted to approximately 0.1% parasitemia in 2 ml dishes (5% hematocrit) and analyzed by FACS as trophozoite stage parasites 1, 3, 5, and 7 days post invasion. After analysis on day 5, parasites were split 1:10 to prevent overgrowth of cultures. For growth analyses over one parasite cycle, highly synchronous ring-stage cultures were diluted to 1% parasitemia in 2 ml dishes and were analyzed by flow cytometry in the subsequent cycle as trophozoite stage parasites.

Flow cytometry-based analysis of growth was performed basically as described previously (Malleret *et al*, 2011). In brief, 20 μl resuspended parasite culture was incubated with dihydroethidium (5 μg/ml) and SYBR green (0.25 × dilution) in medium for 20 minutes at room temperature protected from light. For every sample, 100.000 events were recorded using an ACEA NovoCyte flow cytometer and parasitemia was determined based on SYBR green fluorescence.

For stage quantifications and analysis of schizont morphology, schizont stage parasites were isolated by percoll purification and incubated with uninfected RBCs for three hours to allow rupture and invasion. Parasites were then treated with sorbitol to remove any unruptured schizonts, leading to a 3h synchronous ring stage culture. Next, parasite cultures were diluted to approximately 1% parasitemia in 2 ml dishes (5% hematocrit) and Giemsa-stained blood smears were prepared at 24, 40 and 48 hours post invasion. For analysis of schizont morphology, 1 μM of the egress inhibitor compound 2 was added to schizont stage parasites at 40 hours post invasion. After 8 hours, Giemsa-stained blood smears were prepared and schizont morphology was investigated by light-microscopy.

### Haemozoin crystal motility assay

Schizont stage parasites were isolated by 60% percoll purification and incubated with uninfected RBCs for three hours to allow rupture and invasion. After 3 hours, parasites were treated with sorbitol to remove any unruptured schizonts. Newly invaded rings were then transferred to a 5 mL culture dish (4% haematocrit). Parasites were left untreated or treated with 2.5 mM glucosamine in presence or absence of 100 μM Trolox for 27 hours under standard culture conditions until transfer onto imaging dishes. Treatments with 10 μM diamide in presence or absence of 100 μM Trolox were initiated 24 hours after sorbitol treatment and lasted for 3 hours before parasites were also transferred onto imaging dishes. After the respective treatments, parasites were washed with PBS and trophozoites were isolated by using a Percoll gradient (Heiber & Spielmann, 2014). Enriched trophozoites were then resuspended in 500 μl of complete medium (if necessary with glucosamine, Trolox or diamide) and parasites were allowed to settle on 4 chamber glass bottom dishes (In Vitro Scientific) for 1 hour under standard culture conditions. Imaging dishes were concanavalin A coated beforehand by incubation with 0.5 mg/ml concanavalin A dissolved in H_2_O for 15 minutes at 37°C followed by washing with PBS. Parasites were then imaged using an inverted Leica DFC9000 GT camera and a Leica Plan Apochromat 100x/1.4 oil objective and were kept at 37°C for the duration of the experiment. To generate the videos used for assessing hemozoin crystal motility, one image per second was taken over an imaging period of one minute. For quantifications, video files were imported into ImageJ, where the total number of parasites and the number of parasites with motile hemozoin crystals were quantified.

### Measurement of oxidative stress

For measurement of oxidative stress levels, LCN-Kd and LCN-Con ring stage parasites were treated with 2.5 mM glucosamine or were left untreated. At around 30 hours post-invasion, trophozoite stage parasites were stained with 5 μM CellROX orange (ThermoFisher Scientific) in culture medium for 30 minutes at 37°C. After three washes with PBS, they were directly imaged on a Leica D6B fluorescence microscope as described above. Mean fluorescence intensities and size (area) of individual parasites were determined using the ROI manager of ImageJ.

### Statistical analysis

For statistical analysis of differences between two groups, an unpaired two-tailed students t-test or a ratio-paired two-tailed t-test was used. For statistical analysis of differences between more than two groups, a one-way analysis of variance (ANOVA), followed by a Holm-Sidak multiple-comparison test was performed. P values of <0.05 were considered significant.

## Data availability

All data generated or analyzed during this study are included in this published article and its supplemental material files. The structural data have been deposited in the PDB under the accession code 6TLB. All SAXS data is available on the SASBDB under the accession code SASDH92.

## ACKNOWLEDGEMENTS

We thank Hagai Ginsburg, Murray Junop, Tobias Spielmann and Michael Filarsky for critically reading of the manuscript. Tobias Spielmann is further thanked for providing the aldolase- and SBP1-specific antisera and Mike Blackman is thanked for providing compound 2. Images and time-lapse movies were acquired on microscopes of the CSSB imaging facility.

## AUTHOR CONTRIBUTIONS

Conceived and designed the experiments: PCB, TC, MW, TWG. Performed the experiments: PCB, TC, KL, AZ, CS, BL, LW. Analyzed the data: PCB, TC, KL, AZ, CS, BL, LW, CJ, DIS, DWW, MW, TWG. Performed the phylogenetic analysis: JS. Wrote the paper: PCB, TC, BL, JS, DWW, TWG.

---

**Figure S1.**
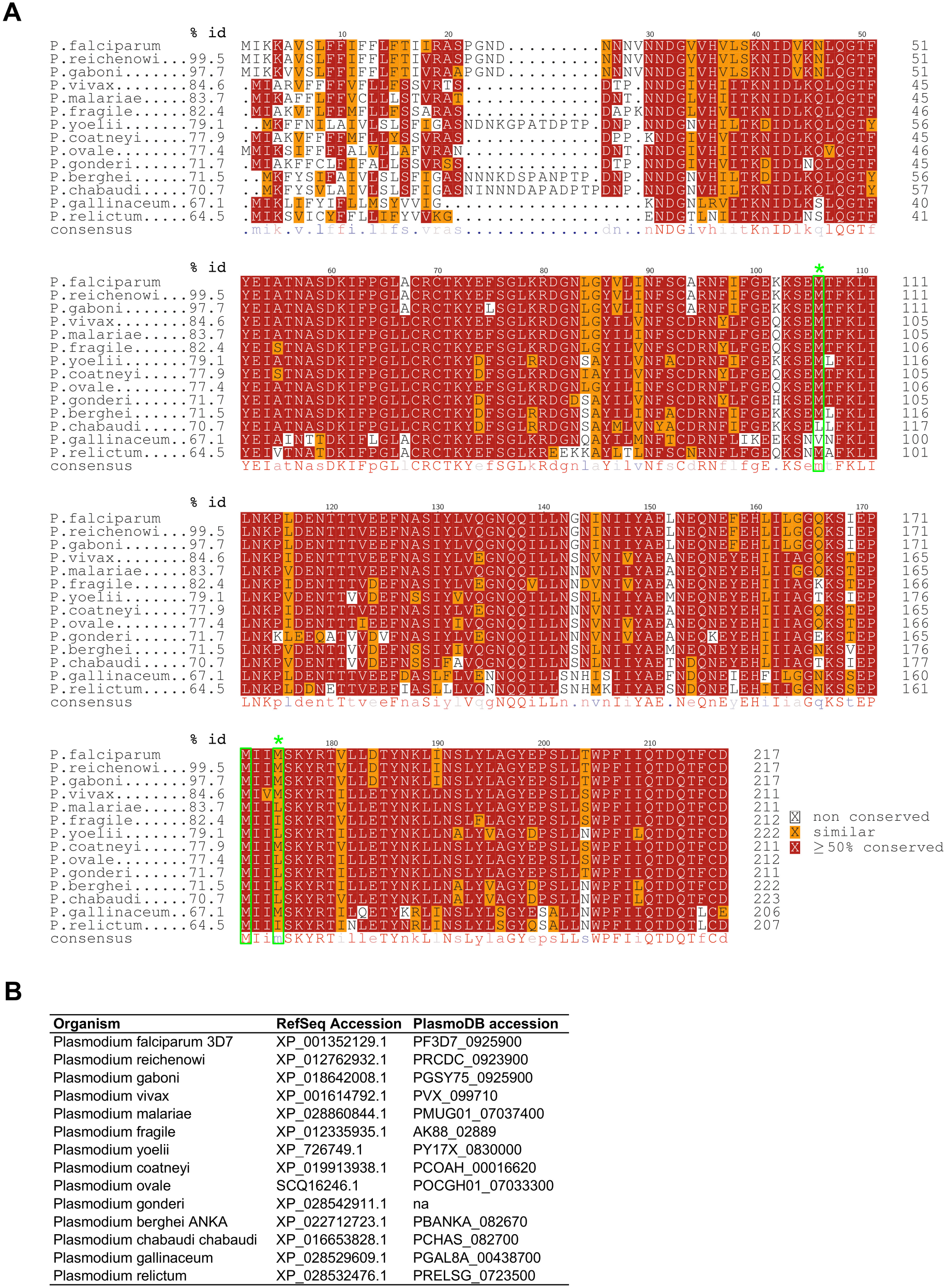
High sequence conservation of lipocalins in different *Plasmodium* species. A) Protein sequence alignment of *Pf*LCN and its orthologues in other *Plasmodium* species and the percentage of sequence identity (% id) compared to *Pf*LCN. Conserved methionine residues are shown in green and those within the *Pf*LCN binding pocket based on its structure are further highlighted with an asterisk. B) Protein accession numbers of proteins used for sequence alignment.

**Figure S2.**
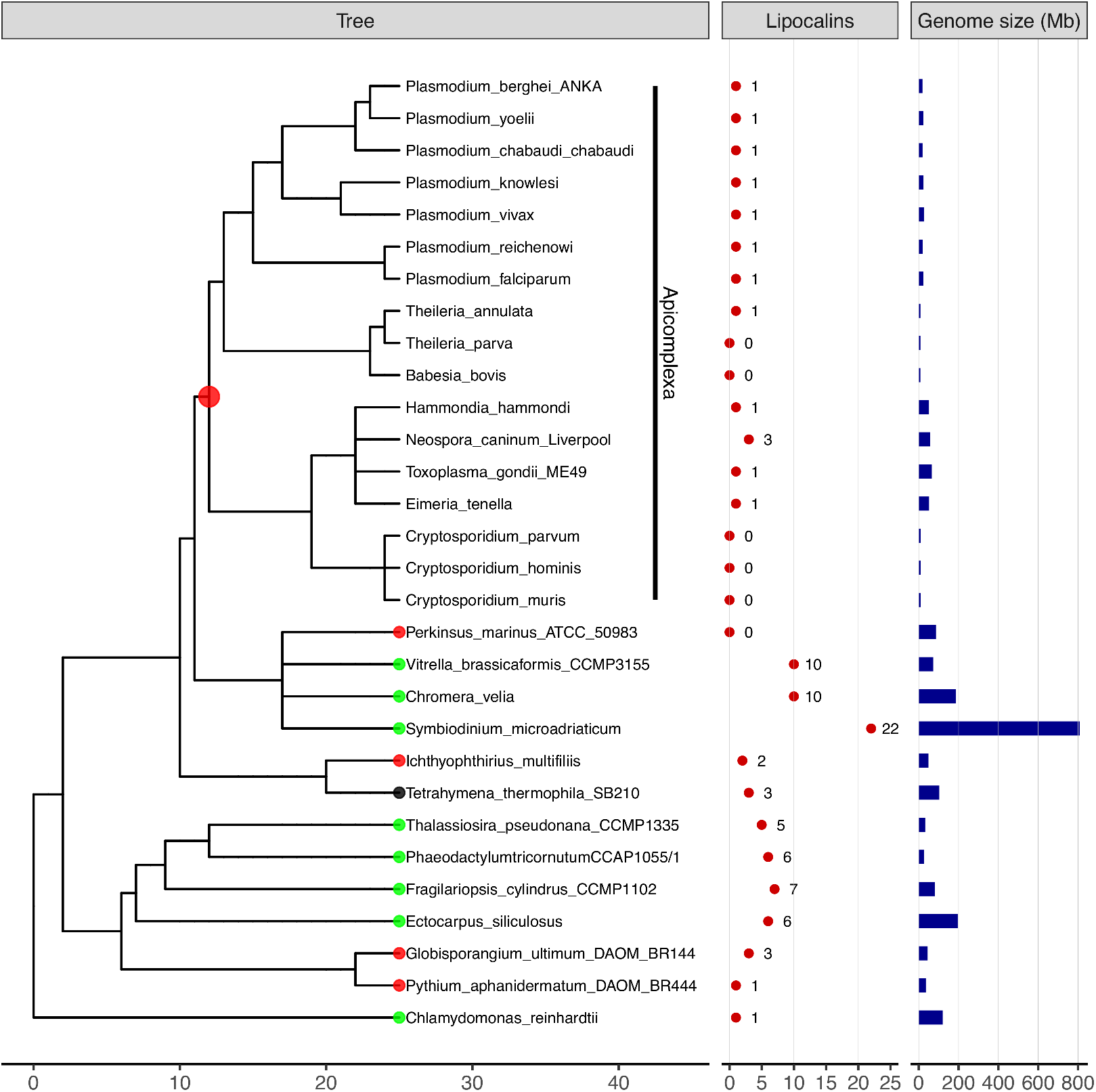
Representation of lipocalin superfamily signatures and genome sizes mapped onto the tree of life of sequenced alveolate taxa including apicomplexan parasites. Phylogenetic tree of alveolate species was derived from open tree of life synthesis v11.4. Apicomplexan species are highlighted with a vertical bar. Nodes and tips of clades and taxa are color-coded by different lifestyles: parasitic (red), photosynthetic (green), and heterotrophic (black). Note that branch length or dates are currently not recorded in the open tree of life synthesis. Protein counts for lipocalin superfamily signatures were derived from InterPro 77.0 and are represented by red points in associated data panel. Genome sizes (Mb) were derived from NCBI genome database and are shown by blue bars in the associated data panel.

**Figure S3.**
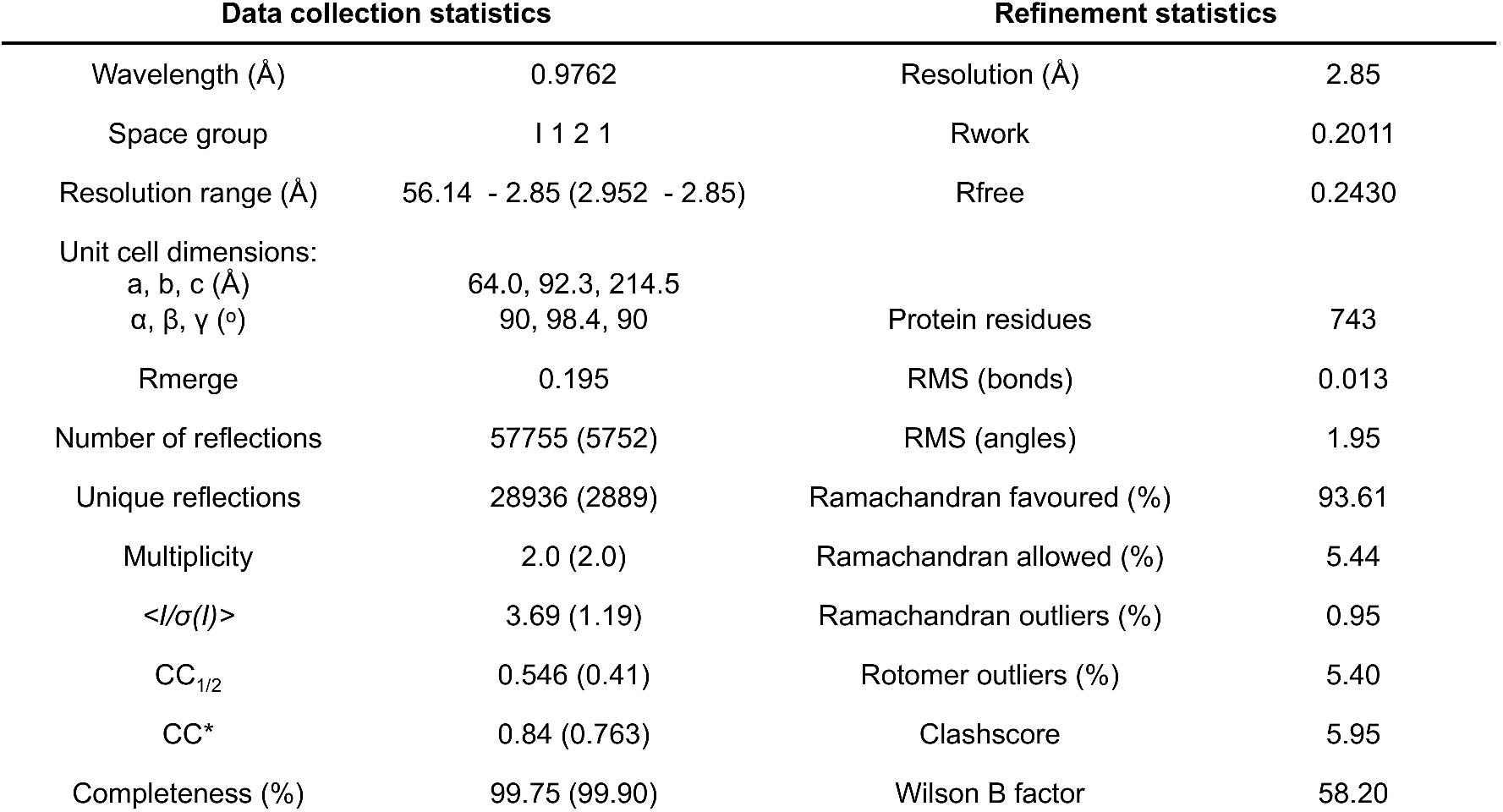
Data collection statistics and refinement statistics of the structural determination of *Pf*LCN. Related to Figure 1.

**Figure S4.**
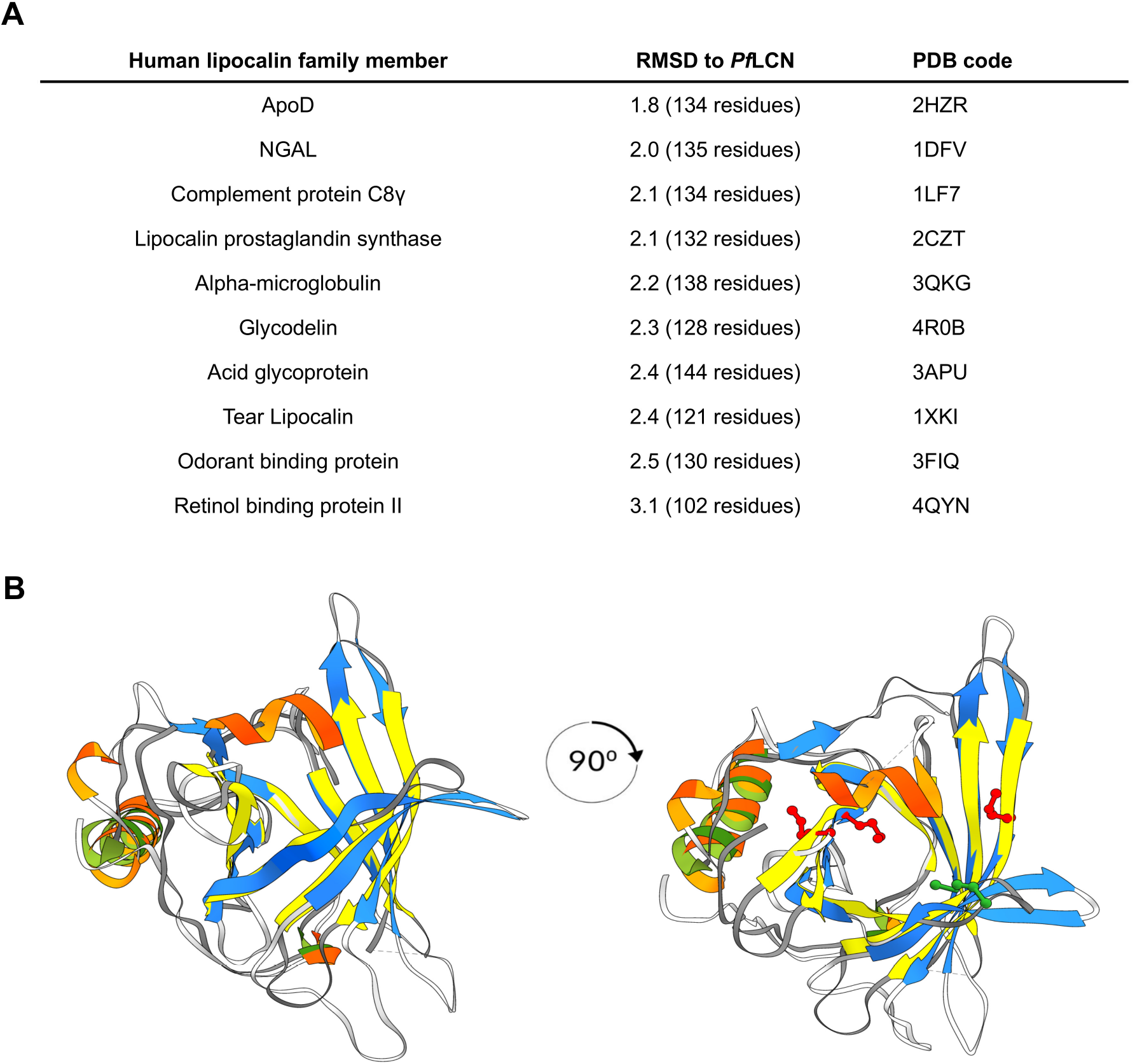
*Pf*LCN shows high structural homology to human ApoD. A) Structural comparisons of *Pf*LCN to the structure of human lipocalins using the program Gesamt. B) Overlay of side-view of *Pf*LCN and ApoD (PDB ID: 2HZR, yellow β-strands and green α-helices). Also depicted is a top-down view of the overlay, with methionine residues highlighted (red on *Pf*LCN: M85, M151, M154; green on ApoD: M93). Related to Figure 1.

**Movie S1. Tetrameric *Pf*LCN.** Related to Figure 1.

**Movie S2. Monomeric *Pf*LCN.** Related to Figure 1.

**Movie S3. Hemozoin crystal motility of untreated LCN-Kd parasites.** One image was taken per second over an imaging period of 1 minute. Scale bar = 5 μm. Related to Figure 5.

**Movie S4. Hemozoin crystal motility of glucosamine-treated LCN-Kd parasites.** One image was taken per second over an imaging period of 1 minute. Scale bar = 5 μm. Related to Figure 5.

**Table S1.**
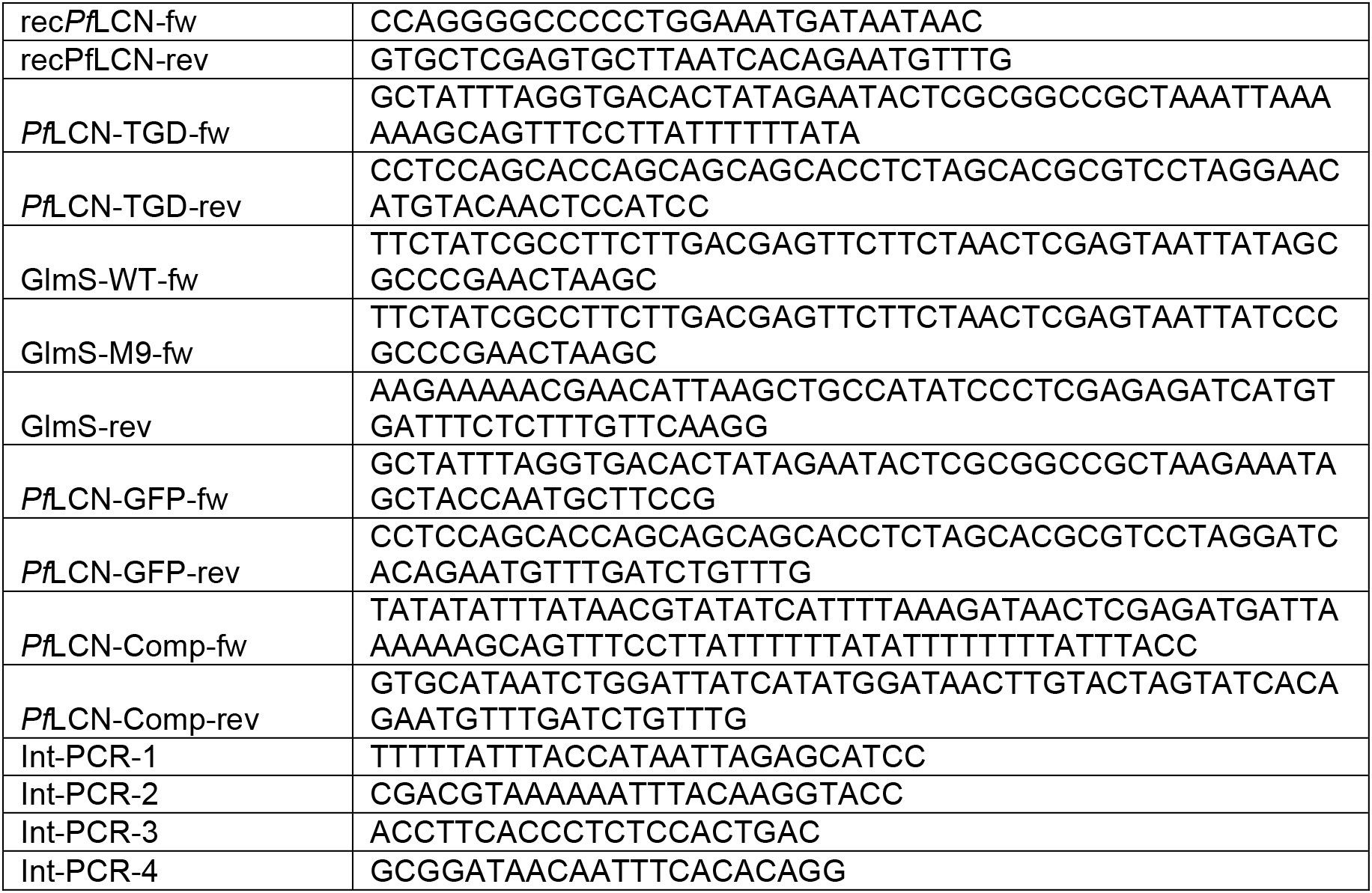
Oligonucleotides used in this study.

